# Deep phenotyping platform for microscopic plant-pathogen interactions

**DOI:** 10.1101/2022.02.17.480879

**Authors:** Stefanie Lück, Dimitar Douchkov

**Author notes:** **Author for correspondence:** Dimitar Douchkov.

## Abstract

- The initial phases of plant-pathogen interactions are critical since they are often decisive for the successful infection. However, these early stages of interaction are typically microscopic, making it challenging to study on a large scale.
- For this reason, using the powdery mildew fungi of cereals as a model, we have developed an automated microscopy pipeline coupled with deep learning-based image analysis for the high-throughput phenotyping of plant-pathogen interactions.
- The system can quantify fungal microcolony count and density, the precise area of the secondary hyphae of each colony, and different morphological parameters. Moreover, the high throughput and sensitivity allow quantifying rare microscopic phenotypes in a large sample size. One of these phenotypes is the cryptic infection of non-adapted pathogens, marking the hidden transition stages of pathogen adaptation and breaking the nonhost barrier. Thus, our tool opens the nonhost resistance phenomenon to genetics and genomics studies.
- We have developed an open-source high-throughput automated microscopy system for phenotyping the initial stages of plant-pathogen interactions, extendable to other microscopic phenotypes and hardware platforms. Furthermore, we have validated the system’s performance in disease resistance screens of genetically diverse barley material and performed Genome-wide associations scans (GWAS), discovering several resistance-associated loci, including conferring nonhost resistance.

## 2 Introduction

Public authorities and society, particularly in Europe, mostly agree about an agroecological transition toward a chemical pesticide-free and GMO-free agriculture. However, this ambitious aim might be challenged by increased outbreaks of new aggressive pathogens promoted by global trade, monocultures, and climatic changes. As high as 40% of global crop production is lost due to pests and diseases, regardless of the extensive use of pesticides (FAO, 2020). Therefore, reduced chemical pesticide use without compensating measures will threaten global food safety to an unacceptable level. One of the most sustainable and environmentally friendly alternatives to chemical pesticides is employing the natural disease resistance of plants. This approach was successfully used in the long history of crop breeding. Still, to meet the new challenges, the plant breeders need to discover new disease resistance sources by digging deep into the genetic diversity stored in the gene banks and germplasm collections worldwide by using more sensitive phenotyping tools capable of discovering quantitative trait loci (QTLs) even with minimal effects and low allele frequency.

The scientific community has identified this need and initiated precise and high-throughput phenotyping tools to establish a new scientific discipline called phenomics. However, most of these efforts were aimed at phenotyping on a larger object level, such as whole plants and canopy, with an insufficient spatial resolution for detailed studies of the typically microscopic plant-pathogen interactions. To contribute to this bottleneck’s alleviation, we started developing a highly automated phenotyping platform to cover the subcellular, tissue, and organ level of phenotyping. The system for organ-level phenotyping on a macroscopic scale called *Macrobot*, and the corresponding software framework (*BluVision Macro*), were published previously (Lück *et al*., 2020; Lueck *et al*., 2020). This article is focused on the high-throughput microscopic system for phenotyping on the cellular and subcellular level, named *BluVision Micro*.

The primary aim of the *BluVision* framework is the phenotyping of plant-pathogen interactions on microscopic and macroscopic levels. As a model for the development was selected, the well-established system of the powdery mildew fungus *Blumeria graminis* as a pathogen of barley and wheat (Panstruga & Dodds, 2009; Spanu & Kamper, 2010; Douchkov *et al*., 2014).

*B. graminis* is the only species of the ascomycete genus *Blumeria*, the order of *Erysiphales*. They are causing powdery mildew diseases on many different grass species. All *Blumeria graminis* are obligate parasites with typically extremely specific host-specialization forms, *called formae speciales (ff*.*spp*.), e.g., *B. graminis* f. sp. *tritici* (wheat powdery mildew, Bgt), and the *B. graminis* f. sp. *hordei* (barley powdery mildew, Bgh) (Wyand and Brown, 2003). Typically the plants are entirely immune against the non-adapted pathogens, e.g., barley is immune to Bgt and wheat to Bgh. However, some plant genotypes may allow microscopic growth of non-adapted pathogens, known as cryptic infection (Romero *et al*., 2018; Bourras *et al*., 2019; Bettgenhaeuser *et al*., 2021). The barley/wheat-powdery mildew model provides several advantages to the researchers: the fungus growth is fast and highly synchronized, the majority of the fungal biomass is located on the leaf surface, with straightforward to observe structures. Furthermore, the fungus interacts only with the uppermost layer of plant leaf cells, the epidermis, via a specialized intracellular feeding organ called a haustorium (Huckelhoven and Panstruga, 2011). This system of reduced complexity provides an excellent environment for studying plant-pathogen interactions on a microscopic scale.

However, full-size and multilevel microscopy images of large objects, such as leaf segments, are typically significant portions of complex data that were only very limitedly accessible with automated image analysis methods until recently. The situation improved dramatically with the coming of age of machine learning (ML) methods that use analytical models to identify patterns and make decisions with minimal human intervention (Mitchell, 1997; Voulodimos *et al*., 2018). There are two main approaches to ML – supervised learning from pre-labeled data (Russell, 2010) and unsupervised learning from unlabeled data (Hinton, 1999). The analysis of images usually includes classification and segmentation steps. The image classification uses features (variables) from images that help classify the objects. The image segmentation assigns labels to the individual pixels, groups them into subgroups (image objects), and subtracts them from the background (Stockman & Shapiro, 2001). Choosing meaningful classification features (feature engineering) (Zheng & Casari, 2018) can be crucial for the success of image analysis. This work compares two main methods - selecting features by human decision (handcrafted features) and automatically extracting features using a convolutional neural network (CNN). CNN can automatically select many features, which leads to more robust prediction models. The downside of the CNNs is the requirement of large training datasets, where predictive models like Random forest (RF) with carefully selected handcrafted features show satisfying results even on small training sets (Lin *et al*., 2020). The optimal approach depends on the specific application and typically would require preliminary testing of different methods.

Here we present the *BluVision Micro* system dedicated to phenotyping the initial stages of plant-pathogen interactions using high-throughput automated microscopy and computer vision methods for localization and quantification of microscopic fungal structures. Unlike the macroscopic systems that typically quantify the disease’s visible symptoms, the *BluVision Micro* delivers precise information about the pathogen behavior, the host’s early response to the pathogen attack, and the fungus’s biomass and growth, virtually eliminating the environment’s effects.

## 3 Related work

The first software development for segmentation and quantifying secondary hyphae of *B. graminis f. sp. hordein (*barley powdery mildew) was the HyphArea Tool (Seiffert & Schweizer, 2005; Baum *et al*., 2011). The software was developed in Python 2. It is based on a histogram-based threshold for hyphae segmentation and a shape descriptor for classifying the regions of interest (ROI).

## 4 Material and methods

### 4.1 Plant and fungal material

Barley cv. Golden Promise and cv. Morex, and wheat cv. Kanzler were grown in 12 cm pots with IPK-soil substrate. The plants were incubated in a plant growth cabinet (Sanyo/Panasonic MLR-352H-PE Versatile Environmental Test Chamber, white LED upgrade; Panasonic Healthcare Co., Ltd.) at controlled conditions (dark period of 8h, light period of 16h, 20°C and 60 RH%) for 7 days or 14 days. The first or the second leaves were harvested and mounted on 1% water agar (Phyto agar, Duchefa, The Netherlands) plates supplemented by 20 mg/L benzimidazole as a senescence inhibitor. The barley leaf segments were inoculated with the Bgh isolate CH4.8, and the wheat leaf segments were inoculated with Bgt isolate FAL92315 at approximately five spores/mm^2^ in an inoculation tower. The fungus was stopped at 36-96 hours after inoculation (hai) by incubating the leave segments in a clearing solution (7 mL 96% Ethanol and 1 mL Acetic acid) for about 48 hours at room temperature. After that, the fungal colonies were stained with Coomassie staining solution (0.3% Coomassie R250, 7.5% (w/v) trichloroacetic acid, and 50% (v/v) methanol) for 5 minutes and then washed several times with water. The prepared samples were mounted on microscope slides with 50 % glycerol to avoid drying the leaves during image acquisition.

The material of the barley core collection of genotypes was grown, collected, and inoculated as described in (Lück *et al*., 2020). In brief, the plants were grown in 24-well seedling trays, ten plants of the same genotype per well, in a climatized greenhouse for 14 days. Leaf fragments from the second leaf were harvested and mounted on standard 4-well microtiter plates, filled with 1% water agar supplemented by 20 mg/L benzimidazole. The leaf fragments were inoculated, incubated, and stained as described above.

### 4.2 Image acquisition and analysis hardware

The microscopy image data was acquired on a commercial *Zeiss AxioScan*.*Z1* high-performance microscopy slide scanner and ZEN 3.0 (blue edition) software (Carl Zeiss AG). The imaging was done in a bright field configuration with a *Hitachi HV-F202SCL* camera (3 CCD 1/1.8” progressive scan color sensor with 1600×1200 effective pixels and 24-bit color depth), 1x camera adapter. As scanning objective typically was used an EC Plan-Neofluar 5x/0.16 M27 with 0.16 NA (numerical aperture) that provides a large depth of field (DoF), which was particularly advantageous for scanning very thick and uneven objects as whole-leaf fragments and helped reduce the Z-stack levels to only five by keeping the most fungal structure focus. The acquired image data was stored in a CZI file container that combines all relevant image and meta information in one file. The image data were analyzed on a Windows 10 Enterprise server with a double *Intel Xeon*™ *E5-2695* processor with 36 physical cores and 512 GB RAM, allowing nearly real-time analysis if required.

The macroscopic image data were acquired six days after infection, as described in (Lück *et al*., 2020). In brief, monochrome images of the leaves illuminated with small-bandwidth isotropic LED light sources with 365 nm (UV), 470 nm (blue), 530 nm (green), and 625 nm (red) peak wavelengths were acquired separately and stored in 16-bit TIFF image files.

### 4.3 Software implementation

The software *BluVision Micro* and all experiments were implemented in *Python 3*.*6* under *Windows 10* operating system. The following free *Python* libraries were used for development: *OpenCV-Python, NumPy, Pandas, Keras, Tensorflow, czifile, skimage, mahotas, joblib* and *Scikit-learn*. Training of the CNN model was done on NVIDIA TITAN X GPU with *Keras* 2.3.1 and *Tensorflow* 2.1.0 backend, and training time of about 20.000 images per hour on an Intel® Core™ i7-9700 CPU 3.00 GHz with 64-Bit Windows 10 operation system.

The software is implemented as a two-step command-line tool with separated image processing and data analysis, allowing curation of the intermediate results without rerunning the entire analysis. In addition, the images processing can be parallelized, depending on the installed computer memory.

### 4.4 Barley Genotyping

Two hundred barley accessions from the barley collection of the Federal *ex-situ* Genebank in Gatersleben, selected for maximized genetic diversity, were genotyped by using whole-genome sequencing (WGS) data from Illumina short-read sequencing with 3x genome coverage (Milner *et al*., 2019), and aligned to the barley MorexV2 reference genome (König *et al*., 2020; Mascher, 2020) (Supplemental Figure S1). A quality filter on 223 387 147 variants was applied with the Plink 2.0 software limiting the missing values to ≤ 0.02 and minor allele frequency (MAF) to ≥0.05. After filtering, 949 174 high-quality variants remained and were used in GWAS analysis.

### 4.5 Best linear unbiased estimator (BLUE)

To obtain robust and unbiased phenotype means for the individual genotypes from the three independent experiment repetitions, we used the Best linear unbiased estimator (BLUE) (Henderson, 1975; Liu *et al*., 2008). BLUE were calculated with the help of the lme4 library for R using the spore inoculation density as fixed factor.

### 4.6 Genome-Wide Association Study

GWAS for the seven traits was conducted with a Factored Spectrally Transformed Linear Mixed Model using a kinship (K) matrix provided by the FaST-LMM program (*fastlmm* 0.5.5) (Lippert *et al*., 2011; Listgarten *et al*., 2012). A suggestive threshold (−log10 P ≥ 6.0) was calculated based on the formula −log10 (1/ number of independent SNPs)(Yang *et al*., 2014) and a significance threshold (−log10 P ≥ 8.0) for the identification of QTLs was calculated by using the Bonferroni correction method (Hommel, 1988).

### 4.7 Phenotype Preprocessing

Six direct phenotypes and one derivative were derived for each genotype from detached leaf samples (Figure 1). The microscopic phenotypes include normalized colony counts at 48 and 96 hours after infection (hai) with the adapted pathogen (Bgh), and 96 hai with the non-adapted fungus (Bgt). In addition, one macroscopic phenotype (infection spread at 168 hai) was included for comparison (Table 1).

**Table 1.** Analyzed phenotypes.

| Phenotype | Phenotyping module | Scale | Pathogen | Time (hai) | Interaction type |
| --- | --- | --- | --- | --- | --- |
| Bgh_48hai_counts | BluVision Micro | Micro | Bgh | 48 | host |
| Bgh_48hai_size | BluVision Micro | Micro | Bgh | 48 | host |
| Bgh_96hai_size | BluVision Micro | Micro | Bgh | 96 | host |
| Bgt_96hai_counts | BluVision Micro | Micro | Bgt | 96 | nonhost |
| Bgt_96hai_counts_bin | BluVision Micro | Micro | Bgt | 96 | nonhost |
| Bgh_48-96hai_slope | BluVision Micro | Micro | Bgh | 48-96 | host |
| Bgh_168hai_area | Macrobot | Macro | Bgh | 168 | host |

**Figure 1.**
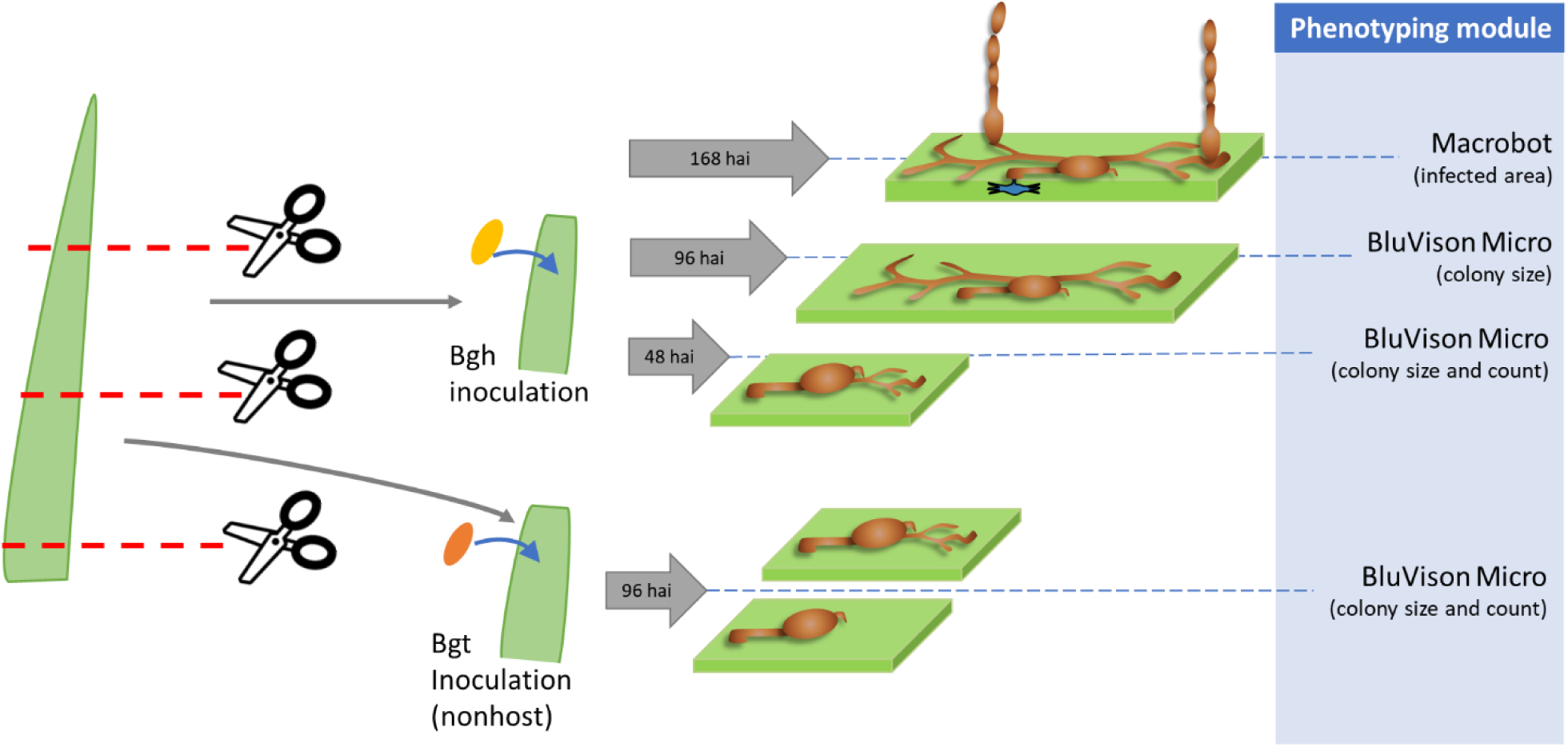
Microscopic and macroscopic phenotypes derived from a single leaf. Up to eight barley plants of the same genotype were grown for 14 days. Two segments from the second leaf of each plant were cut and inoculated with adapted (Bgh) or non-adapted (Bgt) pathogen. Samples for microscopy were collected at 48 and 96 hai, and macroscopic disease rating was done with the remaining leaves at 168 hai.

The colony mean size per leaf at 48 hai and 96 hai Bgh, the colony area was extracted from the segmented images with the OpenCV *contourArea()* function, and the BLUE was calculated from the mean of three experiment repetitions for each barley genotype. The colony sizes at both time points were used to calculate the slope of the growth curve, which was also used as a phenotype in GWAS.

In addition to the quantitative phenotype (normalized colony counts) for the non-adapted pathogen (96 hai Bgt), we also calculated a binary qualitative phenotype using a threshold for the normalized colony count of 0.1. This approach reflects the qualitative nature of the NHR and allows for the identification of major R-genes involved in this complex phenomenon.

The macroscopic infection severity was calculated as the percentage of leaf area covered by the powdery mildew colonies 168 hai using the *BluVision Macro* software (Lueck *et al*., 2020). A mean of up to 8 technical replicates per accession in an experiment was used to calculate the BLUE values from three independed experiemts.

## 5 Results

### 5.1 Image processing

#### 5.1.1 Focus stacking

For finding the optimal focus stacking strategy of the multilevel CZI-images, we have tested five different Z-projection methods included in the *Fiji* distribution package of *ImageJ v1*.*53* - Average intensity (Khamfongkhruea *et al*., 2017), Maximum intensity (Sato *et al*., 1998), Minimum intensity (Hayabuchi *et al*., 2011), Sum slices, Standard deviation and Median (Figure 2).

**Figure 2.**
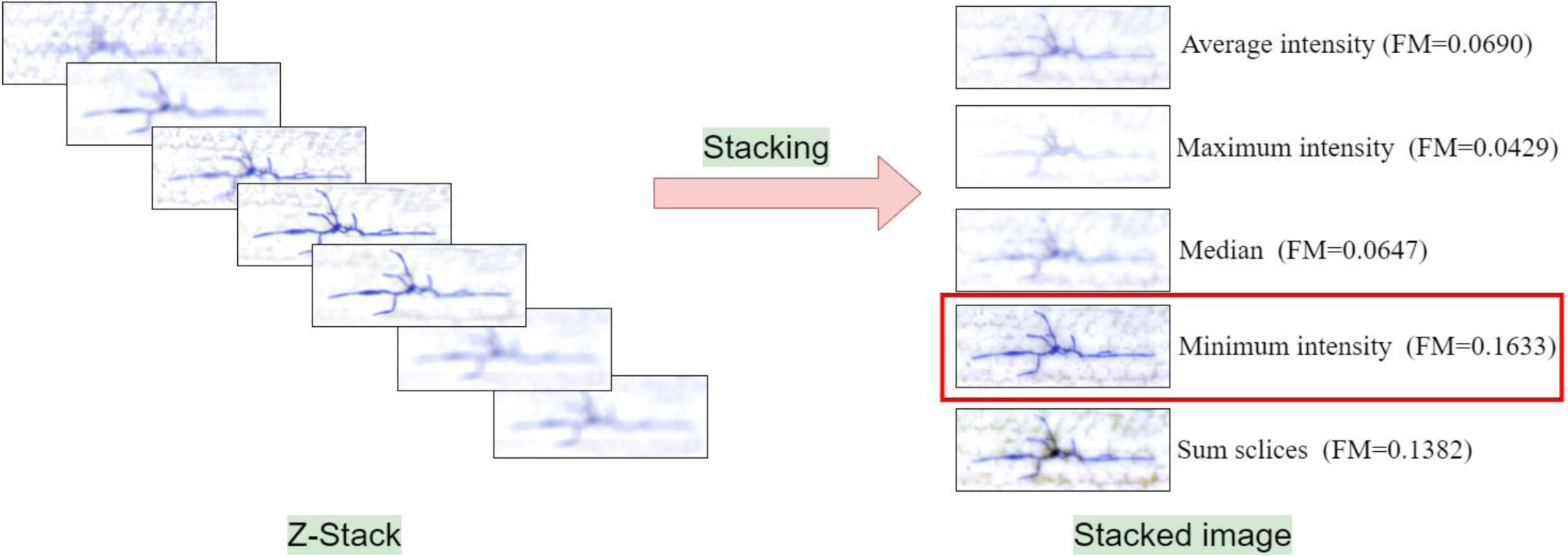
Comparing stacking algorithms. Five stacking algorithms were compared: Average intensity, Maximum intensity, Median, Minimum intensity, Sum slices. The Minimum intensity method achieved the highest quality measure (FM).

Furthermore, for each stacked image, the image Quality Measure (FM) has been computed and compared (Table 2) (De & Masilamani, 2013). The minimum intensity projection method achieved the best FM score in all tested cases and was selected for the image processing pipeline.

**Table 2.** Intensity Z-projection methods compared on two image stacks

| Stack Nr. | Method | FM |
| --- | --- | --- |
| Stack 1 | average | 0.0449 |
| Stack 1 | maximum intensity | 0.0444 |
| Stack 1 | median | 0.0456 |
| Stack 1 | minimum intensity | 0.0825 |
| Stack 1 | sum slices | 0.0691 |
| Stack 2 | average | 0.0690 |
| Stack 2 | maximum intensity | 0.0429 |
| Stack 2 | median | 0.0647 |
| Stack 2 | minimum intensity | 0.1633 |
| Stack 2 | sum slices | 0.1382 |

#### 5.1.2 Colony segmentation

The fungal colony images were extracted and classified in several steps. A significant challenge was to design a reliable pipeline that tolerates staining quality and background variability without losing too many positive objects.

First, the Z-stacked images were segmented to find the putative ROIs. Then, regions of interest were extracted as a bounding box, and the image was classified into a positive or negative class.

Different common color spaces were tested: HSV, L*a*b, YCbCr, XYZ, AC1C2, YUV, I1I2I3 and YQ1Q2, in combination with different thresholding algorithms: Yen’s maximum correlation (Yen *et al*., 1995), Li’s minimum cross-entropy method (Li & Lee, 1993; Li & Tam, 1998; Sezgin & Sankur, 2004), Otsu (Otsu, 1979), Isodata (Ridler & Calvard, 1978), Mean (Glasbey, 1993), Minimum (Prewitt & Mendelsohn, 1966; Glasbey, 1993), Triangle (Zack *et al*., 1977), Canny edge detector (Canny, 1986) (Table 3). Combining the Q2 channel from the YQ1Q2 color space with Yen’s thresholding generated the most reliable results. Using only a single-color channel, we achieved a robust and reliable segmentation method that is insensitive to staining variations and performs well on different sizes of the hyphae (36 to 72 hai).

**Table 3.** Segmentation methods for colony detection (image of 30 000 × 12 000 pixels containing 120 colonies). Comparision of the used software libraries, run-time per image and colony segmentation performance.

| Method | Library | Time(s) | Segmented colonies | Partial segmented colonies |
| --- | --- | --- | --- | --- |
| Canny edge detector | OpenCV | 6.5 | 30 | 109 |
| Global thresholding | OpenCV | 0.3 | 44 | 5 |
| Adaptive Thresholding Mean | OpenCV | 2.0 | 87 | 20 |
| Adaptive Thresholding Gaussian | OpenCV | 3.9 | 86 | 5 |
| Otsu's Binarization | OpenCV | 1.4 | 79 | 16 |
| Li Minimum | scikit-image | 112.0 | 4 | 22 |
| Yen thresholding | scikit-image | 81.0 | 16 | 19 |

A morphological closing operation was applied to the segmented binary images to close the gaps that may lead to partial object extraction. Finally, a Moore-Neighbour tracing algorithm (Weisstein, 2021) was used to extract the contours of the binary image for colony classification.

### 5.2 Machine learning

#### 5.2.1 Training data set

Bgh inoculated barley leaves were Coomassie-stained at 36-72 hai and scanned with the *AxioScan*.*Z1* system. The multilevel images were processed as described above. The putative ROIs were extracted with a bounding box and saved as separate images. The images were manually curated, and about 10 000 ROI containing fungal colonies were selected. Another about 8 000 images without fungal structures but other objects and artifacts were selected as negative training data. Finally, a small training set with 3 200 images per class was extracted from the large training set to study the prediction performance based on the training set size.

Both datasets were split randomly into 75% of the images for training the models and 25% for validation and evaluation. Since the Convolutional Neuronal Network (CNN) approach requires identical dimensions of the training images, they were resized to 150 × 350 pixel, the mean ROI size of the particular data set.

#### 5.2.2 Classification using handcrafted features

Manual selection of features for building a reliable classifier is still a widely used approach that, in some cases, may outperform more sophisticated methods (Lück *et al*., 2020). However, the success of this approach strongly depends on the selection of informative and robust features. In our case, of particular importance was to select color- and scale-invariant features because of the high staining intensity- and colony size variation.

The contours received after the segmentation step were first filtered using geometrical features (Table 4) to reduce artifacts and non-fungal structures.

**Table 4.** Object size parameters for filtering colonies from the artifacts. Minimum and maximum thresholds for colonies are indicated.

| Feature | Minimum pixel values | Minimum pixel values |
| --- | --- | --- |
| Width | 100 | 1400 |
| Height | 100 | 800 |
| Aspect ratio | 0.5 | 10.0 |
| Area | 1000 | 30000 |

Then, five scale- and color-invariant features (Histogram of oriented Gaussians (Dalal & Triggs, 2005), Local binary pattern (Dong-chen & Li, 1990; Wang & He, 1990), Haralick (Haralick *et al*., 1973), Zernike Moments (Tahmasbi *et al*., 2011), Parameter-free threshold adjacency statistics (Coelho *et al*., 2010); Table 5) were extracted with the *mahotas* and *scikit-image* library, and a random forest classifier with 80 trees was trained with the two training sets (3 200 and 10 000 images per class).

**Table 5.** Edge and texture descriptors.

| Name | Abbreviation | Descriptor |
| --- | --- | --- |
| Histogram of oriented Gaussians | HOG | Edge |
| Local binary pattern | LB | Texture |
| Haralick | HA | Texture |
| Zernike Moments | ZM | Shape |
| Parameter-free threshold adjacency statistics | PFTAS | Texture |

Finally, the performance of *Accuracy, Precision*, and *Recall* scores were calculated according to Equation 1 and shown in Tables 6 and 7.

**Table 6.** Random Forest model for image features with 3 200 objects per class. Average of 10 independent trainings.

| Method | Precision | SD | Recall | SD | Accuracy | SD |
| --- | --- | --- | --- | --- | --- | --- |
| HOG | 0.8493 | 0.0097 | 0.8895 | 0.0110 | 0.8634 | 0.0053 |
| LB | 0.9346 | 0.0076 | 0.9547 | 0.0077 | 0.9429 | 0.0048 |
| HA | 0.9075 | 0.0100 | 0.9216 | 0.0071 | 0.9109 | 0.0056 |
| ZM | 0.7816 | 0.0144 | 0.8239 | 0.0075 | 0.7919 | 0.0066 |
| PFTAS | 0.8821 | 0.0070 | 0.9288 | 0.0082 | 0.9000 | 0.0042 |

**Table 7.** Random Forest model for image features with about 10 000 objects per class. Average of 10 independent trainings.

| Method | Precision | SD | Recall | SD | Accuracy | SD |
| --- | --- | --- | --- | --- | --- | --- |
| HOG | 0.8472 | 0.0081 | 0.8893 | 0.0080 | 0.8641 | 0.0059 |
| LB | 0.9346 | 0.0076 | 0.9547 | 0.0077 | 0.9429 | 0.0048 |
| HA | 0.9088 | 0.0057 | 0.9311 | 0.0059 | 0.9186 | 0.0046 |
| ZM | 0.6841 | 0.0116 | 0.7419 | 0.0120 | 0.7018 | 0.0064 |
| PFTAS | 0.8516 | 0.0082 | 0.8830 | 0.0055 | 0.8653 | 0.0056 |

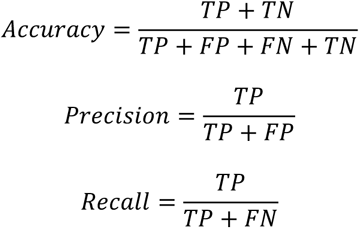

***Equation 1***. *Accuracy, Precision, and Recall scores calculation. TP – true positive, TN – true negative, FP – false positive, FN – false negative (according to the ground thought, see the Validation chapter)*.

#### 5.2.3 Convolutional neural network

We implemented a standard convolutional neural network (Figure 3) with dropout 0.2 and trained two training sets with different sizes (ca. 3 200 and 10 000 images per class) over 25 epochs. We used rectified linear activation function during training, followed by a final SoftMax activation function to receive the probability distribution over the classes. In addition, we used the stochastic gradient descent optimizer with a learning rate of 0.01, batch size of 32, and momentum of 0.9 to allow one training image to pass through the neural network at a time and update the weights for each layer. The final validation accuracy of the model was 97.13% (Figure 4).

**Figure 3.**
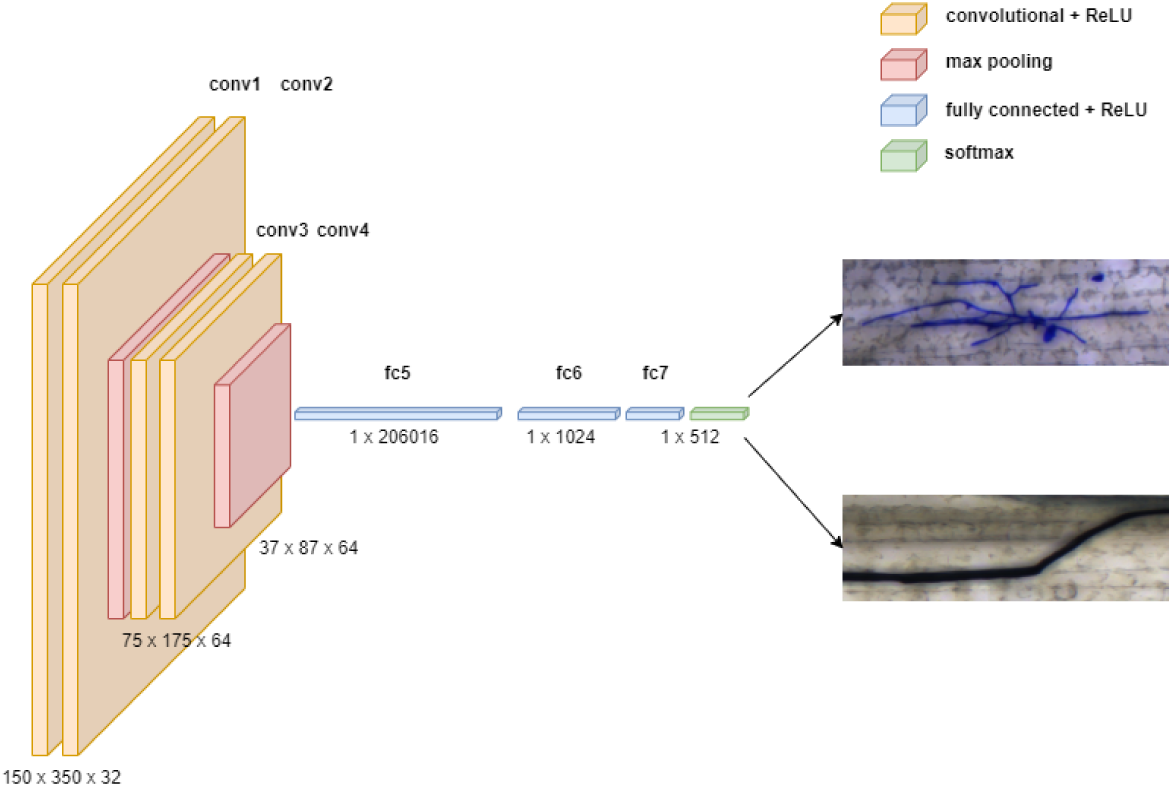
The structure of a convolutional neural network consists of convolutional, pooling, and fully connected layers.

**Figure 4.**
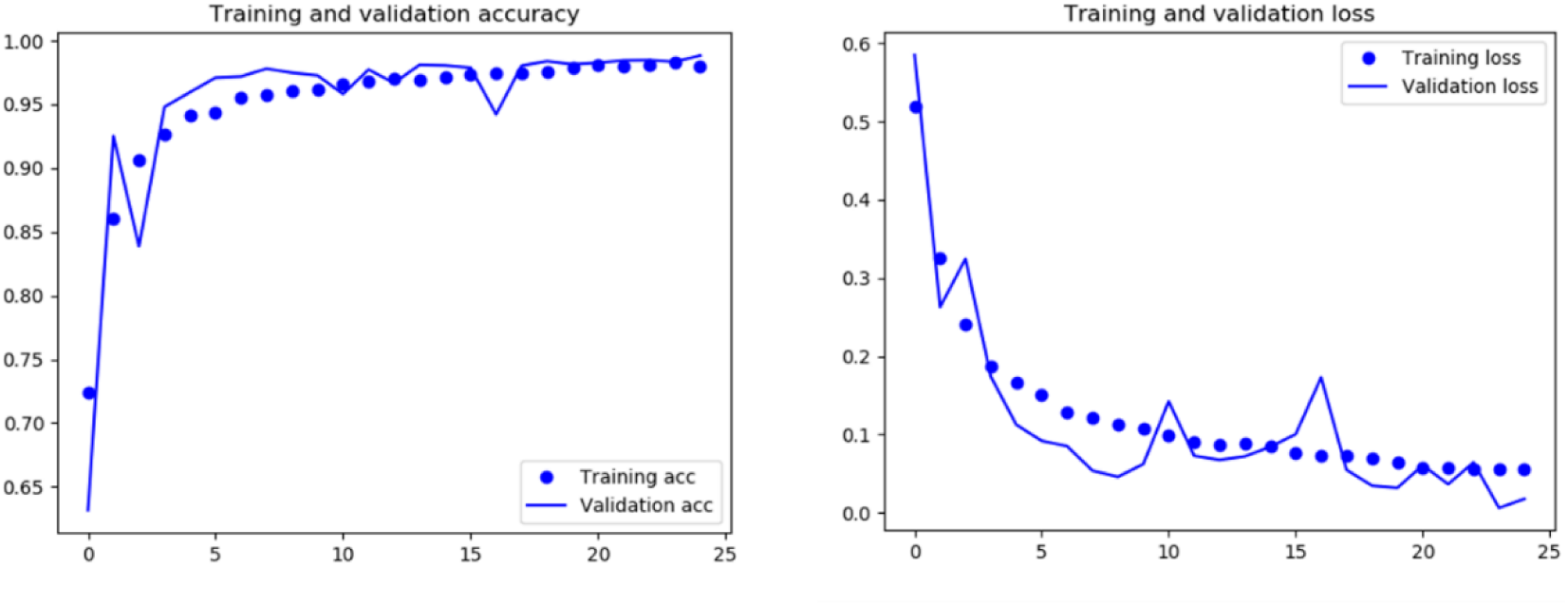
Training and validation accuracy of the model CNN model trained with ca. 10 000 positive images.

### 5.3 Validation

One hundred twenty colonies were labeled manually as ground truth by a domain expert. With the handcrafted feature random forest models trained on 3 200 images per class, the local binary pattern feature reached the highest accuracy and precision (>0.94; Table 6). However, the models ultimately failed on the validation set (False negative > 90%; Table 8). This is usually an indication of model overfitting resulting in a too stringent prediction or a poor capability to deal with new data. This example demonstrates how misleading the theoretical performance metrics can be if used solely without validating the model with new experimental data. Re-testing all previously built models with a new validation data set revealed the *Parameter-free threshold adjacency statistics* (PFTAS) and *haralick* (HA) as best performing (True positives > 88 %, False positives < 10%). Furthermore, a new model based on the combination of both methods significantly improved the accuracy ending up with 91% true positives, 9 % false negatives, and only 1% false positives objects on the validation set (Table 8).

However, increasing the training data size to 10 000 images did not significantly improve the handcrafted feature-based model results, which indicates that the learning curve reached the plateau (Table 7). In contrast, the CNN models usually gain from big data and larger training sets. By using the dataset with 10 000 images, the true positive rate increases by 3.3% to 89.1%, and the false-positive rate decreases to 0.0% (with the prediction accuracy score set to the maximum of 1.0) (Table 8). Loosening the prediction accuracy score to 0.9 helped achieve a high-performance CNN model with over 98% true positive rate and below 3% false-positive rate. In direct comparison, the CNN model shows 10% better accuracy in predicting hyphal objects than the top handcrafted RF-model while keeping the false positive 7% lower (Table 8).

Comparing our best CNN model with a 0.9 prediction score against the HyphArea software, our proposed software improved the true positive prediction by more than 70% and decreased the false positive rate by 10 % (Table 8).

#### 5.3.1 Run-time and parallel processing

Considering the aim for a high-throughput microscopy image analysis, we optimized the algorithm for run-time per image. Besides other improvements, using numerical Python libraries allowing efficient numerical calculations on multi-dimensional arrays, and parallelizing the processes with the *joblib* library (Python) led to a significant speed gain. As a result, *BluVision Micro* performed up to 30 times faster than the previous HyphArea software in analyzing pyramid images of average size 30 000 × 25 000 pixels. On an Intel® Core™ i7-9700 CPU 3.00 GHz with 64-Bit Windows 10 operating system and NVIDIA TITAN X GPU support, the software run time takes about 60 seconds per slide containing two images of size 30 000 × 25 000 pixels, which is 3-5 faster than the image acquisition time, thus allowing real-time analysis.

#### 5.3.2 Feature Visualization

Visualizing the CNN predictions becomes crucial because of the increasing demand for transparency of the artificial intelligence prediction models. However, the availability of visualization options was limited until recently, when several such tools were developed. To examine the *BluVision Micro* CNN model’s prediction and facilitate debugging, we used *Keras Visualization Toolkit (Zhou et al., 2015)* to generate heatmap images to visualize the *Class activation maps* for the fungal structures. The resulting heatmaps correctly represented the area covered by the fungal microcolonies (Figure 5).

**Figure 5.**
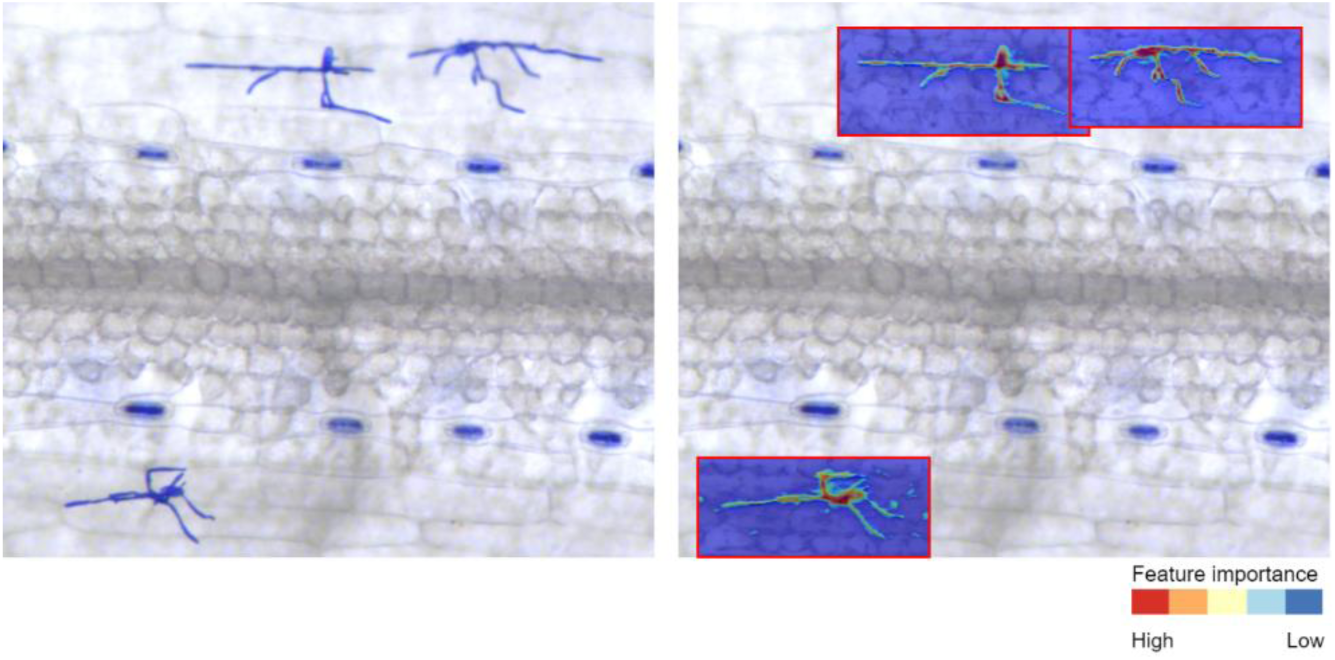
Heatmap visualization of the class activation map for fungal structures. The left image represents the raw image data, and on the right are the regions of interest detected by the software (red border rectangle) with hyphae segmentation. The example clearly shows that the CNN model localizes the fungal colony with high probability (red colors), as the probability in the background drops significantly (blue colors).

### 5.4 Application

#### 5.4.1 Genome-wide association scans (GWAS)

The experiment design (Figure 1) allowed the quantification of multiple phenotypes (Table 1) from a single leaf. The phenotypes cover the response to adapted and non-adapted pathogens on microscopic and macroscopic levels. The precise phenotypic data was combined with the dense SNP data (949 174 quality SNPs) for GWAS for resistance-associated markers.

Since the study aims to provide proof of concept and application examples, the number of tested genotypes was limited to 200, which is on the lower end to detect significant marker-trait associations (MTA) in genetically diverse materials. Nevertheless, we were able to identify eight loci containing MTAs with statistical significance above the suggestive threshold (−log10 P ≥ 6.0) and three loci with MTA above the significance threshold (−log10 P ≥ 8.0). Surprisingly, the novel nonhost resistance phenotypes achieved the highest association peaks leading, besides finding other MTAs, to the re-discovering one of the very few published nonhost resistance QTL (Romero *et al*., 2018) (Figure 9b). All discovered significant MTAs and the genes located in the underlying genomic region are listed in Supplemental Tables MTA_list_[*phenotype_name*] and Gene_list_[*phenotype_name*].

The macroscopic phenotyping (*Bgh_168hai_area*) (Figure 6) suffered from some barley genotypes’ apparent tendency to accelerated senescence in detached leaf assay and formation of physiological necrotic flecks that prevent the spreading of the disease and compromise the phenotyping.

**Figure 6.**
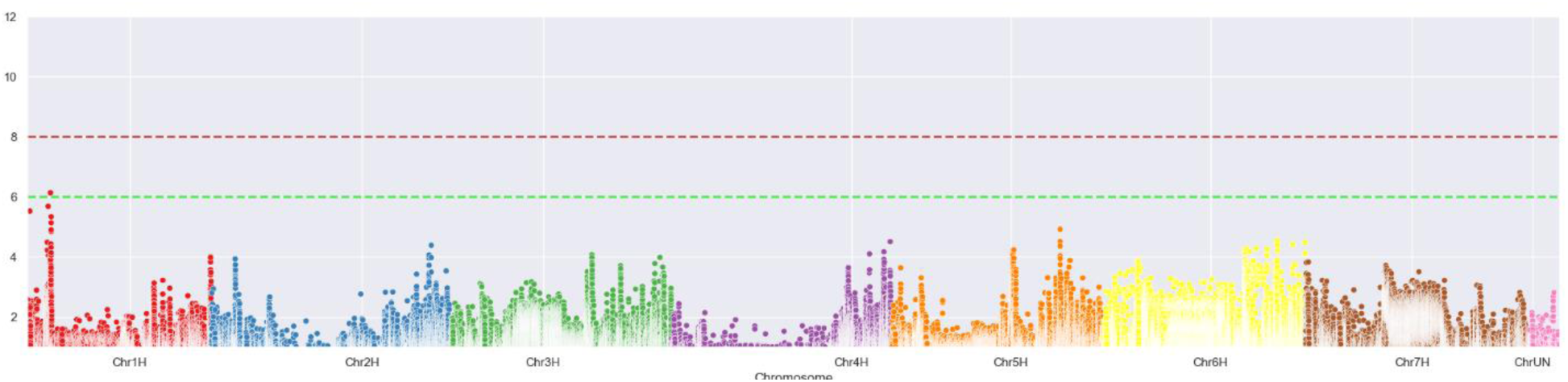
Manhattan plot of the [-log_10_] transformed p-values of the genomic regions associated with the macroscopic phenotype of infected leaf area at 168 hai Bgh (Bgh_168hai_area phenotype). Green dashed line – suggestive threshold, red dashed line – significance threshold.

The colony size-based phenotypes (*Bgh_48hai_size* and *Bgh_96hai_size*) (Figure 7a, respectively 7b) did not deliver significant MTAs. This is not unexpected because a natural resistance in barley based on microscopically-measurable colony growth retardation, to our best knowledge, is not yet described in the literature, not at last because of the lack of screening methods. However, such phenotype likely exists, and a systematic screen of diverse plant genotypes may help discover it.

**Figure 7.**
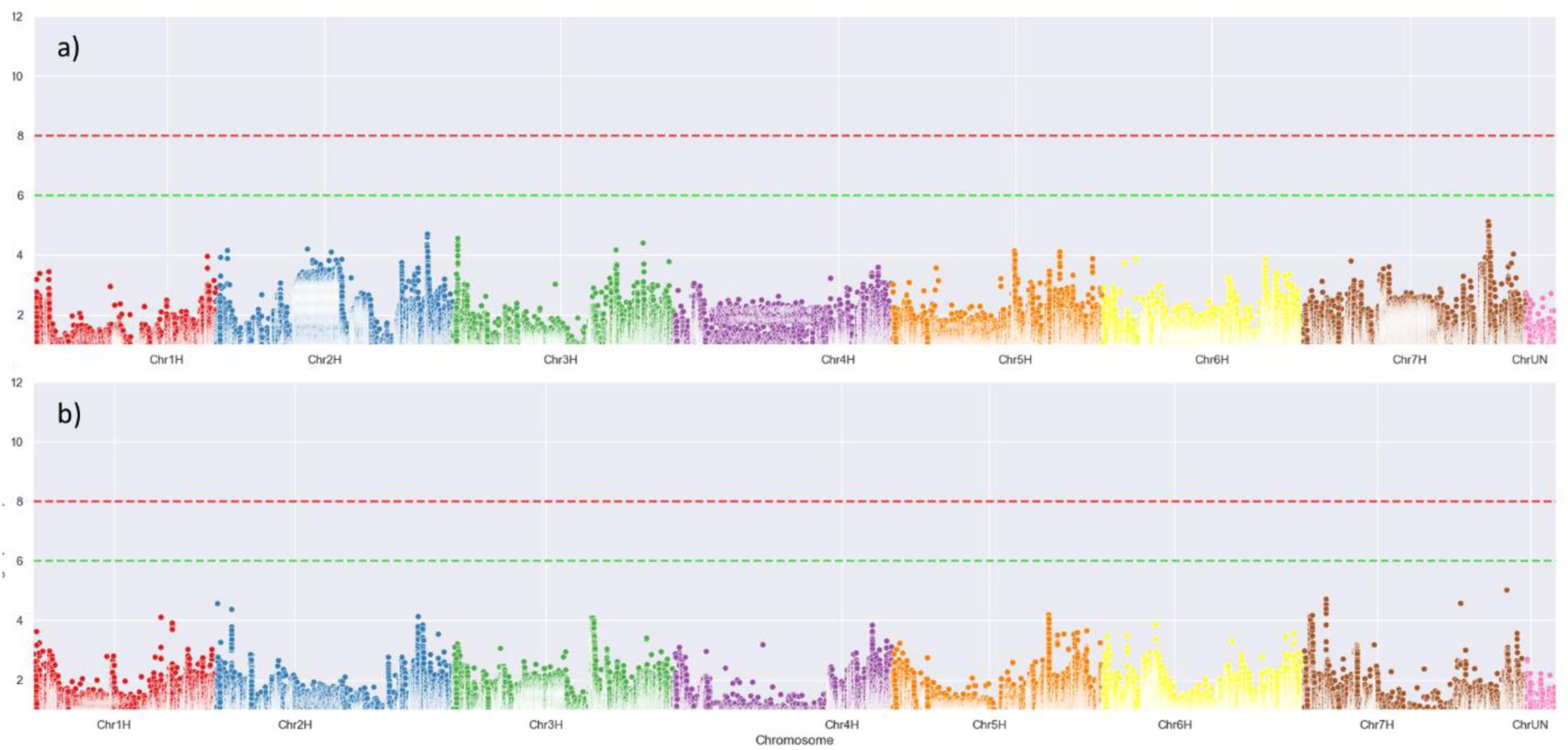
Manhattan plot of the [-log_10_] transformed p-values of the genomic regions associated with colony size-based phenotypes a) Bgh colony size at 48 hai (Bgh_48hai_size), b) Bgh colony size at 96 hai (Bgh_96hai_size). Green dashed line – suggestive threshold, red dashed line – significance threshold.

Also, as expected, the colony counts delivered some significant MTAs (Figure 8) since the penetration resistance against powdery mildew fungus, which effectively reduces the number of successful infection events, is widespread in barley. However, the MTAs reached only the suggestive threshold, not the significance threshold, which was relatively high because of the large number of SNP included in the analysis (∼1 000 000).

**Figure 8.**
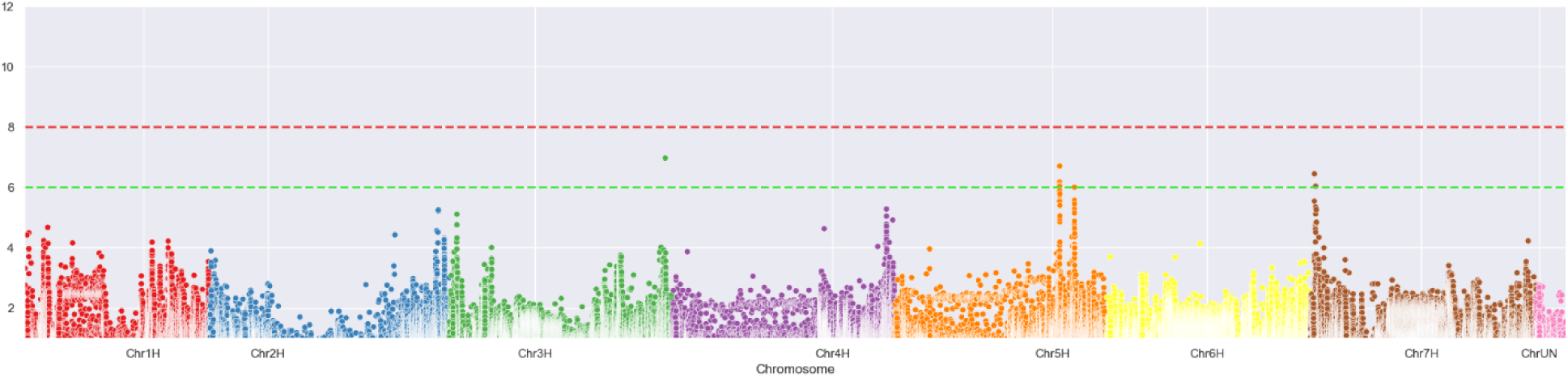
Manhattan plot of the [-log_10_] transformed p-values of the genomic regions associated with normalized Bgh colony counts at 48 hai (Bgh_48hai_counts). Green dashed line – suggestive threshold, red dashed line – significance threshold.

The high sensitivity and performance of the system allowed approaching an exciting novel phenotype – quantifying the rare cryptic infection of non-adapted pathogens, which allowed to monitor the hidden transitions stages of pathogen adaptation to new hosts. At the same time, assigning quantitative phenotype to the honhost resistance opens this most valuable type of resistance to associations studies for the discovery of genes and loci associated with this most valuable type of resistance (Figure 9).

**Figure 9.**
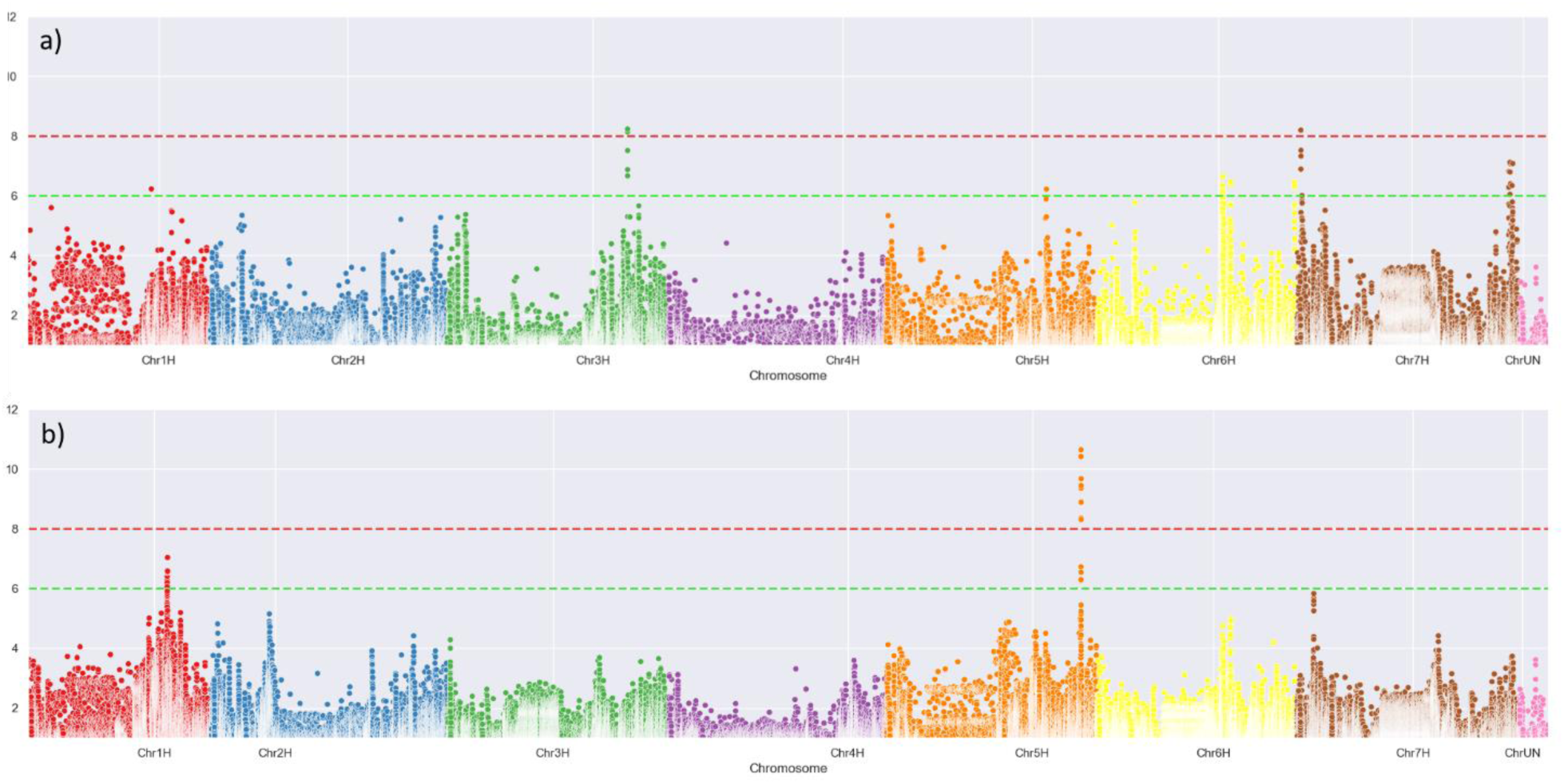
Manhattan plot of the [-log_10_] transformed p-values of the genomic regions associated with (a) normalized Bgt colony counts (Bgt_96hai_counts), and (b) - binarized susceptibility phenotype at 96 hai (Bgt_96hai_counts_bin). Green dashed line – suggestive threshold, red dashed line – significance threshold.

Surprisingly, this novel phenotype delivered the most significant MTAs, indicating the involvement of major-effect genes. Furthermore, the MTA with the absolute most significant p-value in the entire experiment pointed precisely to the peak marker position found by (Romero *et al*., 2018) and probably conferred by one or both of the Receptor-like kinases located in this region.

#### 5.4.2 Pathogen growth curves

The *BluVision Micro* platform provides the possibility to measure precisely, and in high-throughput, the area of the secondary hyphae of the powdery mildew colonies. This opens new phenotyping options, hardly possible with manual microscopy. For instance, measuring the colony size at a specific time point after inoculation may reveal plant defense mechanisms that rely on retarding the pathogen growth, e.g., cutting the nutrient support for the fungus or late activation of cell death mechanisms. Furthermore, acquiring colony size data on multiple time points will allow for building growth curves for the pathogen (Figure 10).

**Figure 10.**
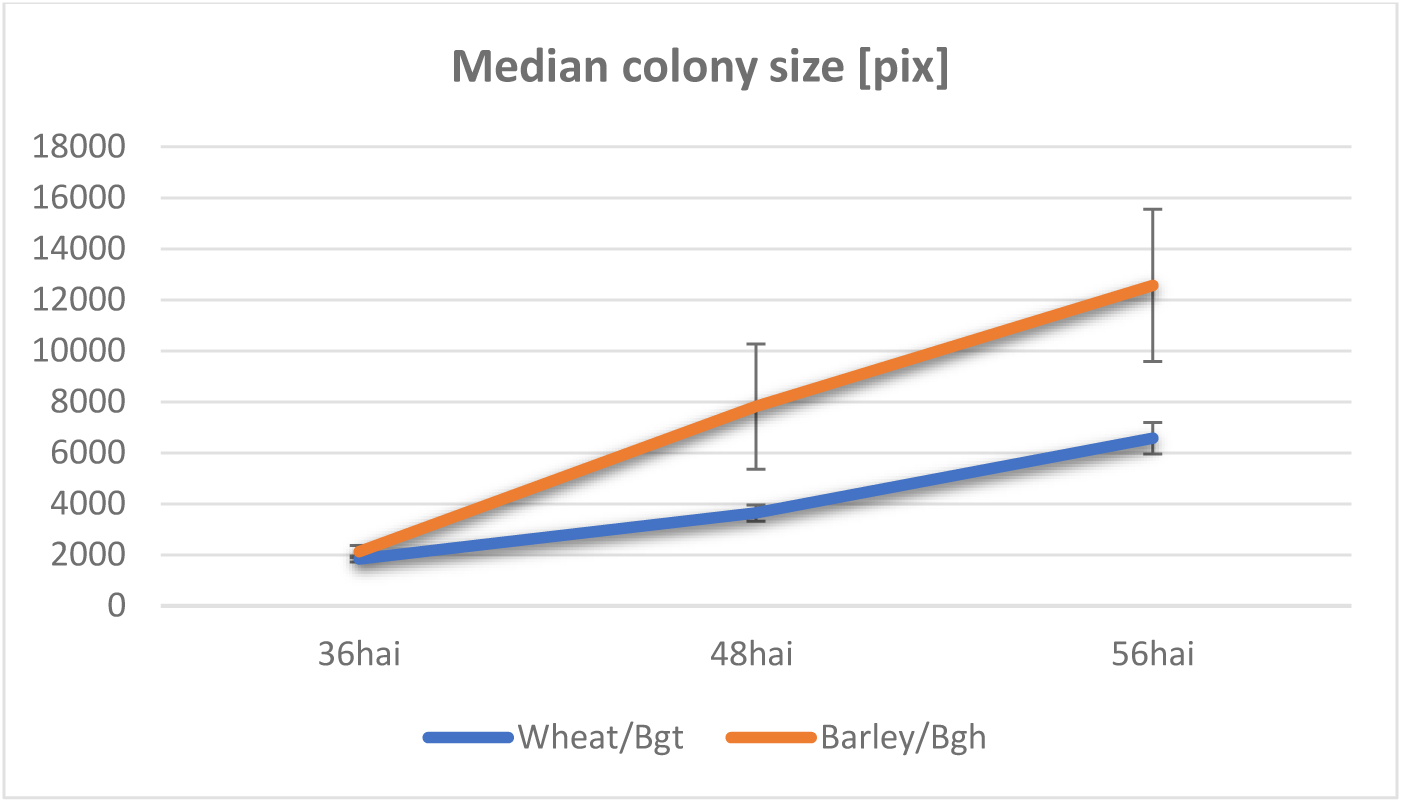
Growth curve of two adapted powdery mildew species on wheat and barley, respectively.

We used the median Bgh colony sizes at 48 and 96 hai on the 200 barley genotypes to build genotype-specific growth slopes and used them as a phenotype in GWAS. As for the direct colony size phenotypes, none of the MTAs reached even the suggestive threshold with the derivative one. Nevertheless, this novel phenotyping method may reveal plant resistance that works by affecting the growth rate of the pathogen. Also, it can be a valuable tool in comparing the fitness of different pathogen races.

**Figure 11.**
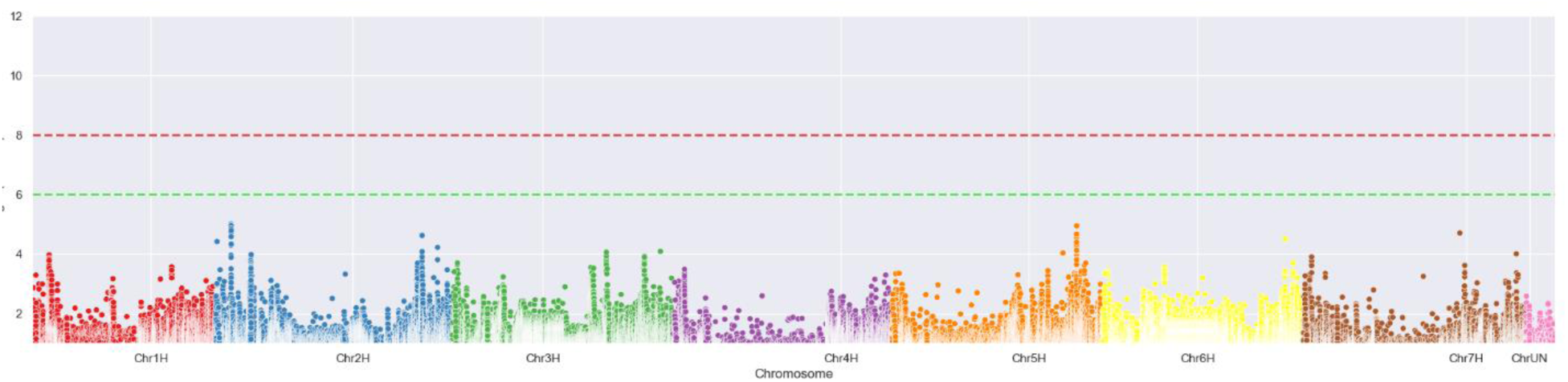
Manhattan plot of the [-log_10_] transformed p-values of the genomic regions associated with the slope of the growth curve of Bgh at 48-96 hai (Bgh_48-96hai_clope). Green dashed line – suggestive threshold, red dashed line – significance threshold.

## 6 Discussion

The need for automated microscopic phenotyping of plant-pathogen interactions became apparent with increasing the number of available genetic and genomics resources and the pursuit of finding novel genes putatively involved in the complex phenomenon of disease resistance. HyphArea was the first software implementation to detect and quantify secondary hyphae of *B. graminis* on barley and wheat. The tool pioneered establishing a high-throughput platform for plant-pathogen interaction phenotyping on a microscopic level allowed access to novel phenotypes such as quantification of the area of fungal secondary hyphae. However, the high sensitivity and specificity levels of the HyphArea Tool demonstrated in (Seiffert & Schweizer, 2005; Baum *et al*., 2011) was often difficult to reach due to the vaiability of the sample properties and quality.

Besides the image analysis, the extended use of the HyphArea revealed issues with the handling and processing of the raw data. The acquired image data were exported as individual camera frames (tiles) and stored in separate TIFF files. This step simplifies the image data processing and avoids using proprietary file formats but results in a massive expansion of the file number (>10^6^ files for a large screen), thus approaching the limits of the used hardware and software. Finally, the long run time of the HyphArea renders it less appropriate for high-throughput phenotyping screenings.

Benefiting from the accumulated experience and using newer high-throughput automated microscopy and software techniques, we have developed a completely new system for microscopy-based phenotyping. We decided to opt for a modular, machine learning-based software that works directly with different image data types, including complex pyramid files and multimodal images, and it is easily adaptable and extendable with modules for additional phenotypes.

Different machine-learning (ML) approaches were tested and evaluated. Handcrafted features-based ML models, if chosen correctly, can provide acceptable performance in cases where only small (< 5 000 images per class) training sets are available. However, using more training data for the handcrafted features approach does not further increase the performance, showing that we have reached the methods’ limits in this case. For higher accuracy and larger training sets (> 5 000 images per class), we recommend using a CNN, which significant advantage is extracting the probability for each class and using it as a parameter for predictions.

The newly developed *BluVision Micro* system can derive precise microscopic phenotyping information for different large-scale studies, such as screening of Genebank material, crossing populations, mutant collections, breeding material, and others, at both host and pathogen sides. In this study, we have used the system to screen 200 genetically highly diverse barley genotypes for interaction phenotypes with adapted and non-adapted powdery mildew fungi. The system was confirmed to deliver accurate, sensitive, and reproducible results. We have used them to scan for marker-trait associations in the barley genome, discover several novel loci, and confirm already known. Noticeably, we were able to re-discover one of the first published nonhost-resistance QTL, described by (Romero *et al*., 2018), which confirms the system’s applicability for studies aiming to discover genes involved in this precious but hardly accessible trait – the nonhost resistance. Furthermore, the system allows high-throughput studies of previously extremely laborious phenotypes, such as precise colony area and scoring pre- and post-haustorial defense reactions. By using other (not yet published) dedicated modules, the BluVision platform can also detect the presence of fungal haustoria in reporter gene (GUS) expressing cells, thus enabling high-throughput transfection assays for disease resistance-related genes. The open-source software system allows the development of specific modules for other microscopic phenotypes. The framework is hardware-independent and adaptable to different commercial imaging systems based on the Digital Imaging and Communications in Medicine (DICOM) standard, such as Zeiss Axionscan and Leica Aperio systems.

Thus, we have developed an open-source, extendable, high-throughput automated system for analyses of microscopic phenotypes. Furthermore, we have validated the system’s performance in disease resistance screens of genetically diverse barley material and demonstrated that the phenotypic data could be used for Genome-wide associations scans (GWAS), discovering several resistance-associated loci, including conferring nonhost resistance.

## 7 Acknowledgments

This work was initiated and actively supported by Patrick Schweizer, who passed away in March 2018. The authors dedicate this work to his memory. Further on, we would like to acknowledge the following colleagues from IPK Gatersleben: Nils Stein, Martin Marcher, Murukarthick Jayakodi for the genotypic data for the 200 barley accessions; Andreas Börner for providing single-seed-descent seed material for the experiments; André Fessel and Gabriele Brantin for the technical help, Jens Bauernfeind for the IT support, and Daniel Arend for the data submission. The work was supported by IPK Gatersleben and the German Ministry of Education and Research (BMBF) with grants FKZ 031A053 (DPPN), FKZ 031B0184 (GeneBank 2.0) and FKZ 031B0196 (PrimedPlant).

## 8 Author Contribution

DD designed the research and conducted the biological experiments. SL performed the image analysis, computer model development, and GWAS. SL and DD analysed the data and wrote the manuscript.

## 9 Data Availability

*BluVision Micro* ships with Attribution-NonCommercial 4.0 International license (CC BY-NC 4.0). The open-source code is accessible at https://github.com/snowformatics/BluVisionMicro. Image training sets are available at the electronic Data Archive Library (e!DAL) under http://dx.doi.org/10.5447/ipk/2022/1 (Lueck, 2022).

