## Supplementa Figure S1 for "Deep phenotyping platform for microscopic plant-pathogen interactions"

Supplemental Figure S1. Principal component analysis of the two hundred barley accessions (https://bridge.ipk-gatersleben.de/#pcaplot)

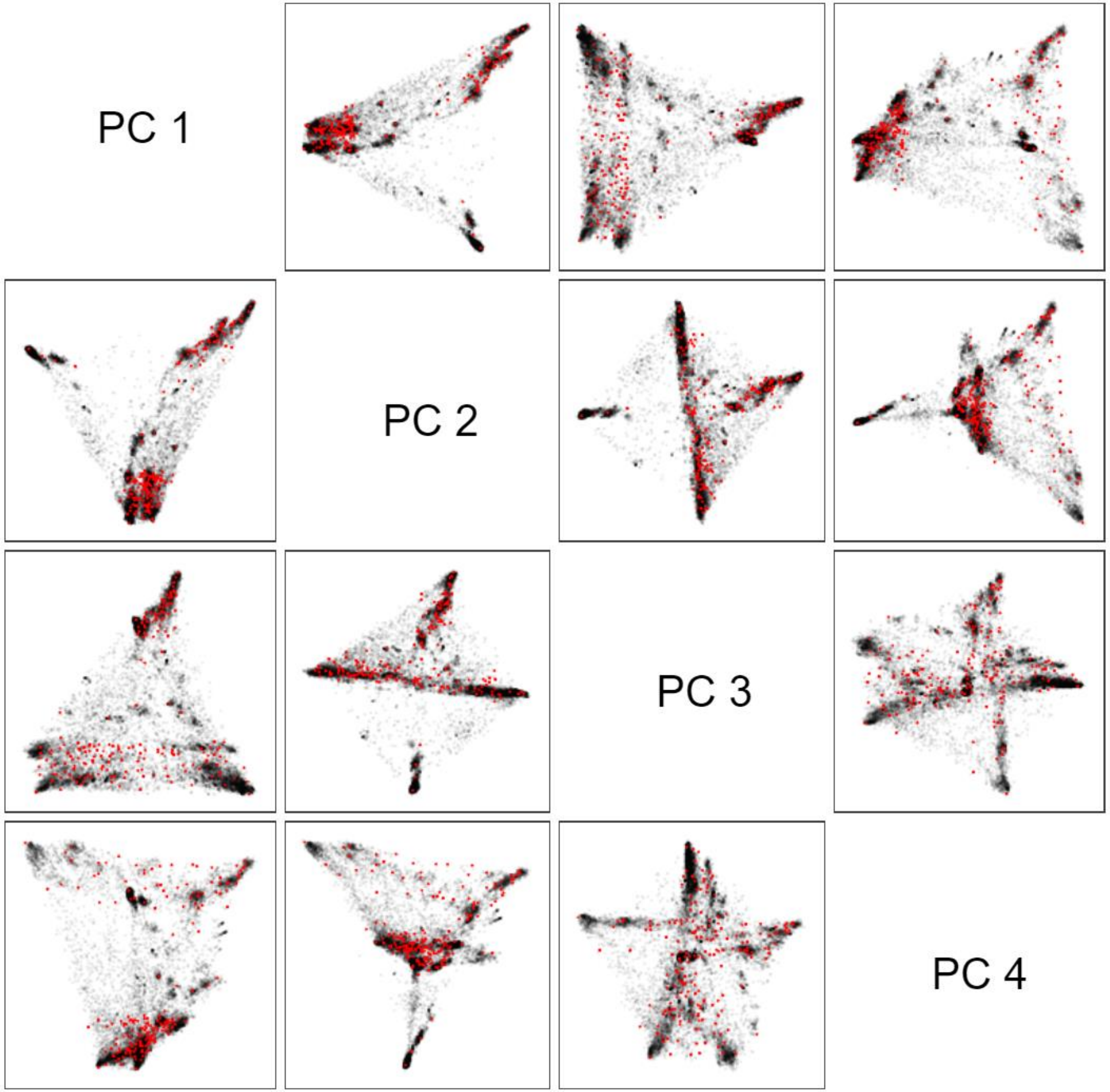
